# Tilt Aftereffect Spreads Across the Visual Field

**DOI:** 10.1101/2022.06.21.496978

**Authors:** Busra Tugce Gurbuz, Huseyin Boyaci

## Abstract

The tilt aftereffect (TAE) is observed when adaptation to a tilted contour alters the perceived tilt of a subsequently presented contour. Thus far, the TAE has been treated as a local aftereffect observed only at the location of the adapter. Whether and how TAE spreads to other locations in the visual field has not been systematically studied. Here we sought an answer to this question by measuring TAE magnitudes at locations including but not limited to the adapter location. The adapter was a tilted grating presented at the same peripheral location throughout an experimental session. In a single trial, participants indicated the perceived tilt of a test grating presented after the adapter at one of fifteen locations in the same hemifield as the adapter. We found non-zero TAE magnitudes in all locations tested, showing that the effect spreads across the tested visual hemifield. Next, to establish a link between neuronal activity and the behavioral results, and to predict possible origins of the spread, we built a model based on known characteristics of neuronal responses in the visual cortex. Simulation results showed that the model can successfully capture the pattern of the behavioral results. Furthermore, the pattern of the optimized receptive field sizes suggests that mid-level visual areas, for example V4, could be critically involved in TAE and its spread across the visual field.

## Introduction

Perceived orientation of a two-dimensional contour is strongly affected by the observer’s state of adaptation. For example, sustained exposure to an oriented contour may change the perceived orientation of a subsequently presented contour, a phenomenon known as the tilt aftereffect (TAE) (Gibson & Radner, 1937). Besides orientation, similar adaptation effects are observed in other stimulus dimensions, such as motion (Anstis, Verstraten, & Mather, 1998), color (McCollough, 1965), and size (Blakemore & Sutton, 1969).

Previous studies have commonly treated TAE as a local effect, and investigated the behavioral and neuronal responses with test contours presented at the same location as the adapter. Physiological studies done in this way revealed two main changes in neuronal responses as a result of adaptation: (1) relative fatigue or suppression in response amplitudes of neurons tuned closer to the adapter orientation (Blakemore & Campbell, 1969), and (2) shift in neurons’ preference away from the adapter orientation (Dragoi, Sharma, & Sur, 2000). These two mechanisms lead to different perceptual outcomes. The former leads to a repulsive shift in population response, and causes a perception away from the adapter orientation. In contrast, the latter leads to a population response attracted towards the adapter and causes a perception closer to the adapter orientation. Models combining the two mechanisms were able to predict TAE more successfully compared to those based on only a single mechanism (Jin, Dragoi, Sur, & Seung, 2005). Accordingly, it was argued that both mechanisms play a role in TAE; the suppression of neuronal responses constitutes the repulsive effect observed in TAE, whereas shifts in neuronal preferences weaken the effect and reduce perceptual errors. Figure 1 depicts the simultaneous effect of these two mechanisms.

**Figure 1:**
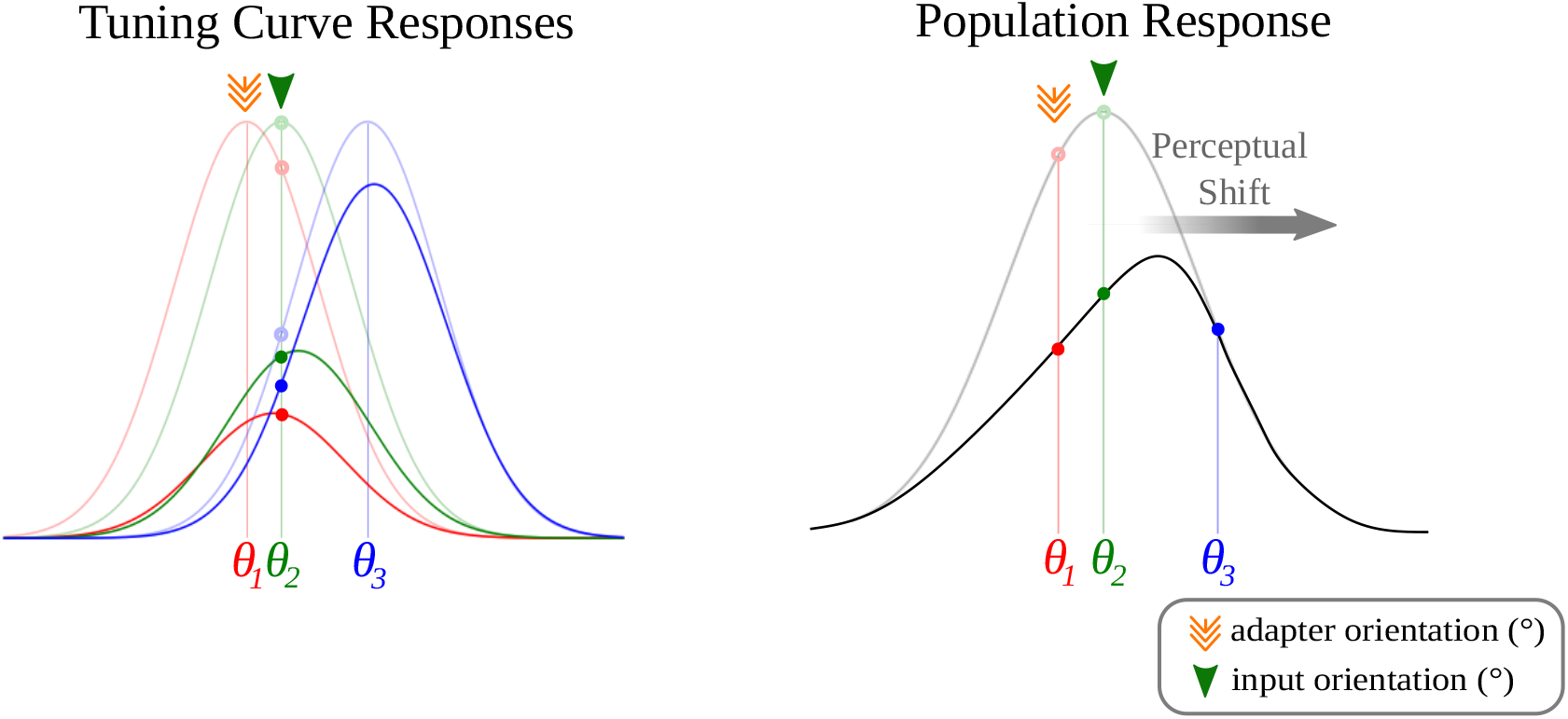
The suppression and shift models of orientation adaptation. In both panels the thin lines represent unadapted neural responses, thick lines represent the responses after adaptation. The left panel shows examples of hypothesized responses of neurons with different tuning curves peaking at several orientations (*θ*_1_, *θ*_2_, *θ*_3_). The right panel shows the population responses obtained from the activity of those neurons. Here we show how the responses change after adapting to an orientation of *θ*_1_. On the left panel, unfilled circles represent the responses of neurons to an orientation of *θ*_2_ before adaptation. The population response obtained with this pattern of activity leads to a peak at *θ*_2_ as shown on the right panel. As a result of adaptation to *θ*_1_, the responses of neurons that are tuned closer to *θ*_1_ have stronger amplitude suppression than those tuned to further orientations. Moreover, their orientation preference shift more strongly away from the adapter. The net result of this suppression and shift lead to an overall repulsive shift away from the adapter orientation in the population response. This shift in neural responses can in principle predict the perceived shift in orientation, and the tilt aftereffect (TAE). Note that this figure is formed by hypothetical shift and suppression values for demonstration purposes.

There may be benefits of this type of adaptation aftereffects for the visual system. For instance, it could serve as a gain control mechanism to maximize visual processing efficiency by increasing the salience of the novel visual input (McDermott, Malkoc, Mulligan, & Webster, 2010). Naturally, one might expect that these benefits would not be tied to the adapter location to allow efficient processing across the visual field. In general, however, TAE and other perceptual aftereffects are commonly considered localized with few exceptions (see for example Altan and Boyaci (2020) for the spread of size adaptation aftereffect across the visual field). How the perceived tilt of a contour is affected by an adapter located at another location has not been investigated systematically before. Understanding this is critical since the visual information from different spatial locations is integrated to form our perceptions and actions in natural visual environments.

In this study, we investigate how adaptation to a tilted Gabor patch at one location in the visual hemifield impact the perceived tilt at other locations. For this purpose, we used two main experimental procedures: a selective-adaptation procedure to a peripheral adapter at a fixed location in the visual hemifield and a testing procedure to test the TAE magnitude at fifteen test locations within the same visual hemifield, including the adapter location. Furthermore, to understand how neuronal populations integrate information from different spatial locations and hypothesize about the possible visual cortical areas involved in this integration, we developed and tested a computational model using the aforementioned suppression and shift mechanisms of adaptation.

## Experiment

### Materials and Methods

#### Participants

19 right-handed participants (8 Female, 11 Male; age range: 19-27, *M* = 21.1; *SD* = 2.1) with normal or corrected vision took part in the experiment. Participants gave their written informed consent before the experiment. Ethical approval for the experiment was obtained from Bilkent University Human Ethics Committee.

#### Stimuli and Apparatus

Stimuli were presented on a 30-inch NEC MultiSync LCD monitor (60 Hz refresh rate and 1920×1200 pixels resolution) in a dark room. A chin-rest was used to stabilize participants’ heads.

Two main stimuli were used in the experiment: adapter and test stimuli, both of which were Gabor patches generated in Python (Version 3.7, available at http://www.python.org) using the PsychoPy library (Version 3.0.1.) (Peirce et al., 2019) in a way that would elicit the maximum TAE magnitude based on previous studies in literature (Harris & Calvert, 1989). All patches had a diameter of 3° (2.7° visible stimuli), 1.0 Michelson contrast, and 1.44 *c/deg* spatial frequency. Adapter patch was always tilted clockwise, with an orientation of 105° (left-handed coordinate system, where 90° is the vertical orientation), whereas two 1-up-1-down adaptive staircases per test location determined the test patch orientation. Each staircase contained 25 trials, and their starting orientations were 80° and 105°. Within each staircase, orientation of the test patch in a trial was adjusted by the experimental program based on participant’s responses in previous trials. The step size of the adjustment was 2.0° at the beginning of the procedure, and it was reduced to 1.0° after the first reversal of participant’s orientation judgment, and to 0.5° after the second reversal, and was kept at that level until the end of the trials in that staircase (Nishida, Motoyoshi, Andersen, & Shimojo, 2003).

Participants sat 65 cm away from the screen and fixated a central fixation point (FP). During an experimental session the stimuli were presented at only one visual hemifield. Left and right visual hemifield tests were counterbalanced across sessions and participants. The adapter patch flickered with a square-wave modulation at 10 Hz to prevent afterimages and was always presented at 10.5° eccentricity. The test patch was presented at one of the fifteen possible test locations, randomly chosen in a trial, in the same visual hemifield, including the adapter location as shown in Figure 2. Three rows of the test locations were named relative to the adapter as Center (*C*; same row), Down (*D*; 4° lower row), and Up (*U*; 4° upper row). Five columns of the test locations were named relative to the location of the adapter patch as Center (*C*; same column as the adapter), Near1 and Near2 (*N1, N2*; closer to FP than the adapter by 4° and 8°, respectively), and Further1 and Further2 (*F1, F2*; further from FP than the adapter by 4° and 8°, respectively). Consequently, names of the test gratings were formed by combining their row and column names (e.g., *C*_*N*1_; center row, Near1 column). Because of symmetry, the same test patch names were used for both visual hemifields.

**Figure 2:**
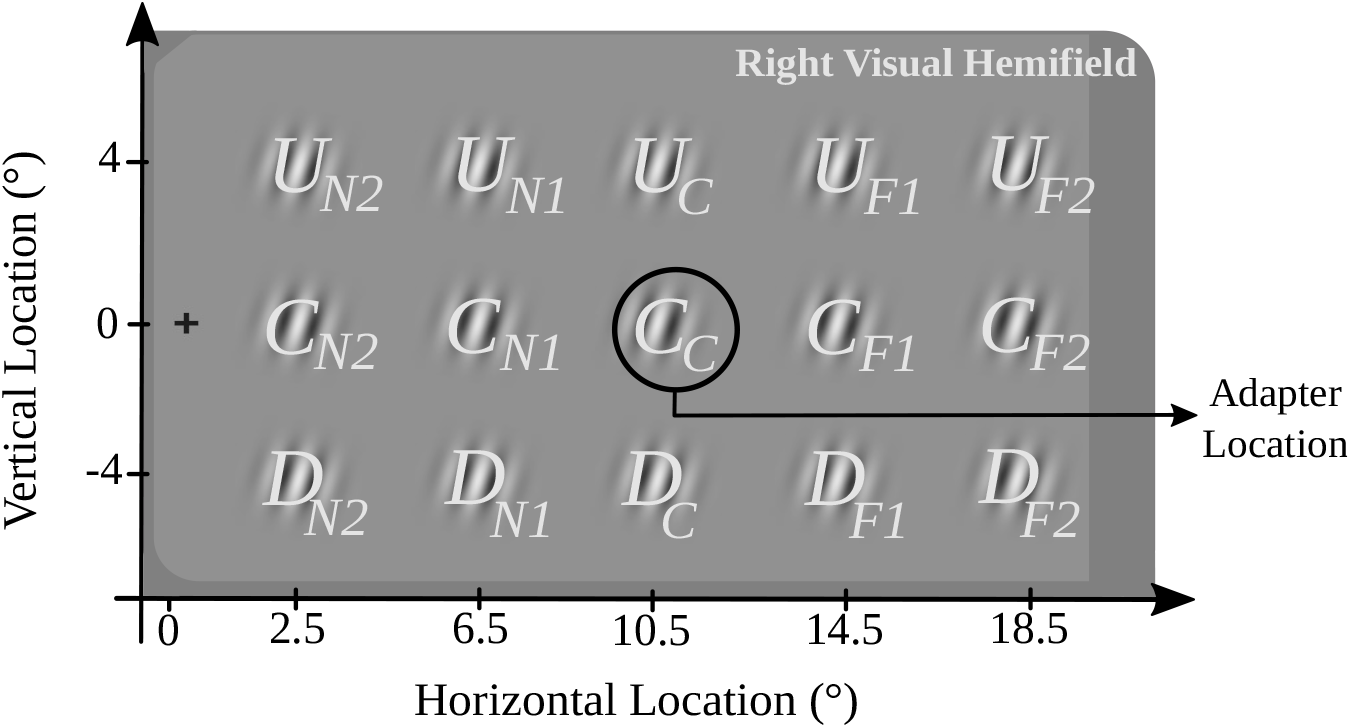
Naming convention, and locations of the adapter and test patches in the right visual hemifield. Left visual hemifield locations are mirror symmetric of these. The cross is the central fixation point (FP), and the solid circle marks the location of the adapter grating. The first letter of the location names indicates its row relative to the adapter (C: same row; D: 4° lower; U: 4° upper). The second name refers to the column relative to the adapter (C: same column as the adapter; N1 and N2: nearer to the FP than the adapter by 4° and 8°, respectively; F1 and F2: further from FP than the adapter by 4° and 8°, respectively.)

#### Procedure and Data Analysis

The experimental procedure was adapted from Altan and Boyaci (2020) and the code for the experimental procedure is publicly available (see Data and Code Availability Statement section). Participants started an experimental block by pressing the space bar on a keyboard after reading the experiment instructions on the screen. They completed the no-adaptation blocks before the adaptation blocks. Figure 3 shows the experimental design of both conditions. For adaptation condition, an experimental trial started with 5000 ms adaptation to adapter grating and it was followed by a 200 ms blank screen and 100 ms test grating presentation. Then, a 2000 ms blank screen was presented where participants indicated the perceived tilt of the test grating by pressing left (for counterclockwise perceived tilt) or right (for clockwise perceived tilt) arrow key button on a keyboard. No-adaptation condition was identical to the adaptation condition except there was no adapter before the test. The experiment contained 1500 trials in total (2 experimental conditions [Adaptation and No-adaptation] x 2 adaptive staircases x 25 trials x 15 test locations). Participants were debriefed after the experiment.

**Figure 3:**
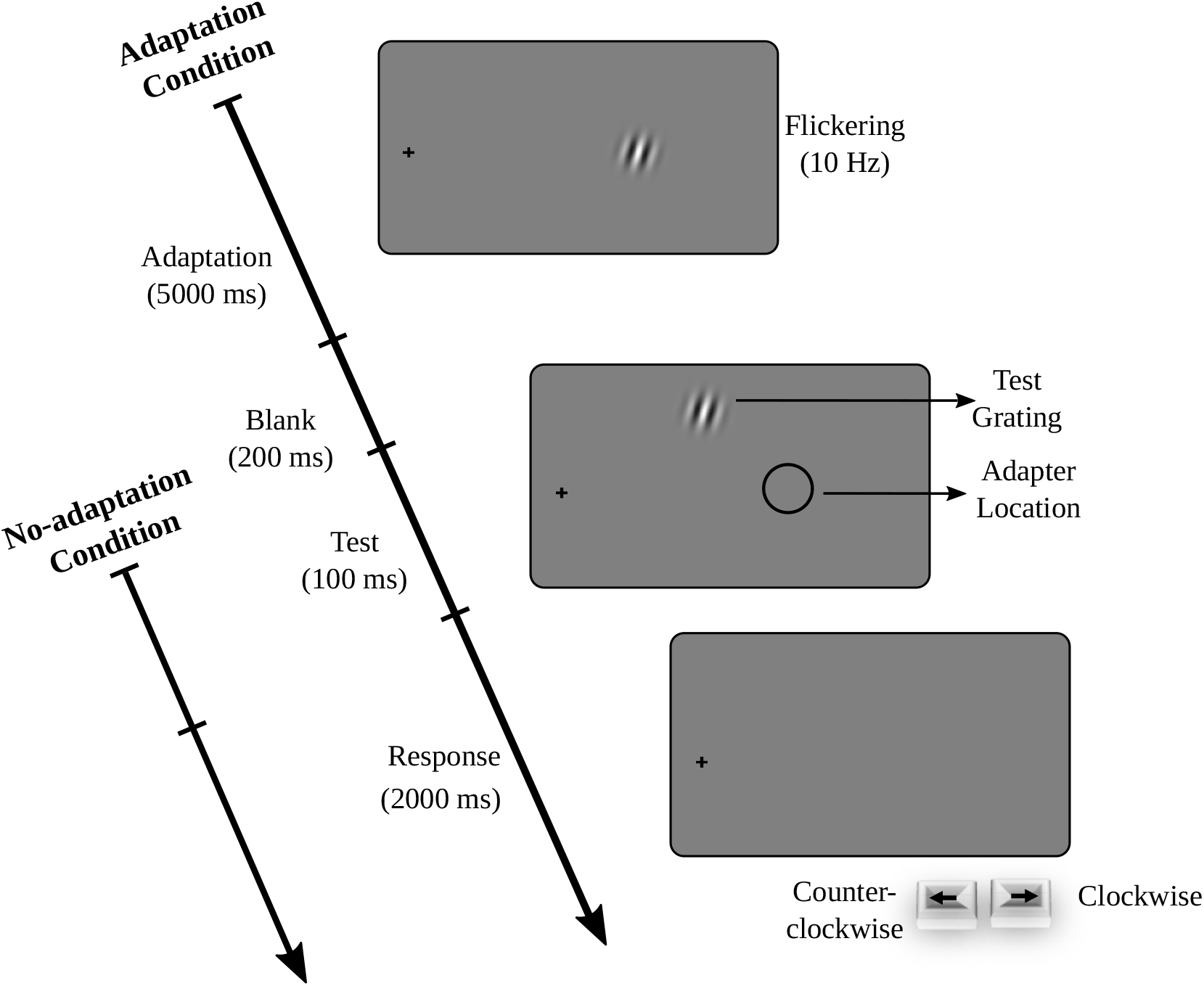
Experimental design. Adaptation blocks started with a 5000 ms adaptation to a patch that was always presented at the same location (*C*_*C*_) and tilted clockwise. After a 200 ms blank screen, the test grating was presented at a randomly chosen position among fifteen possible positions (see Figure 2) for 100 ms. During the following 2000 ms blank screen, participants indicated the perceived tilt of the test grating by pressing the right (i.e., clockwise) or left (i.e., counter-clockwise) arrow key on a keyboard. The no-adaptation condition was identical to the adaptation condition, except for the adaptation phase.

We calculated the point of subjective verticality (PSV) (i.e., the angle at which the participant would report the perceived tilt as clockwise 50% of the time) by fitting a logistic regression function using Psignifit 4 MATLAB Toolbox (Schütt, Harmeling, Macke, & Wichmann, 2016). For each participant, 30 PSV values were calculated (2 experimental conditions x 15 test locations) using an equal asymptote method of Psignifit. Next, the TAE magnitude was calculated as

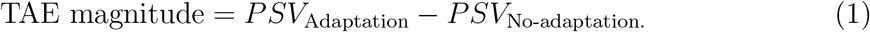

A positive TAE magnitude implies a counter-clockwise ‘repulsion’ away from the adapter orientation, a negative value implies a clockwise ‘attraction’ towards the adapter orientation.

Statistical analyses were performed on the calculated TAE magnitudes using the JASP software (JASP Team, 2020). First, to answer our research question, we performed a pre-planned two-tailed one sample t-test to test whether the TAE magnitude is significantly different from zero at each test location (TAE magnitude ≠ 0) using the pooled data from both visual hemifields. FDR correction was performed to correct for multiple comparisions. Additionally, we performed a mixed ANOVA with post-hoc pairwise t-tests to test the effects of the row, column, and visual hemifield (i.e., within and between-subject factors) on the TAE magnitude.

## Results

Figure 4 top panel shows the TAE magnitudes for all test locations as a heatmap. Because the TAE magnitudes did not systematically vary between the hemifields, the results are averaged and represented on the right visual hemifield (no main effect of visual hemifield in between-subject mixed ANOVA, *F* (1, 17) = 2.16, *p* = 0.16). This map clearly shows that the TAE is not limited to the adapter location, and it spreads across the visual hemifield. Pre-planned one sample t-tests revealed that this spread of the TAE was significant at all test locations (FDR corrected *ps <* 0.05) except *D*_*N*2_ (FDR corrected *p* = 0.17).

**Figure 4:**
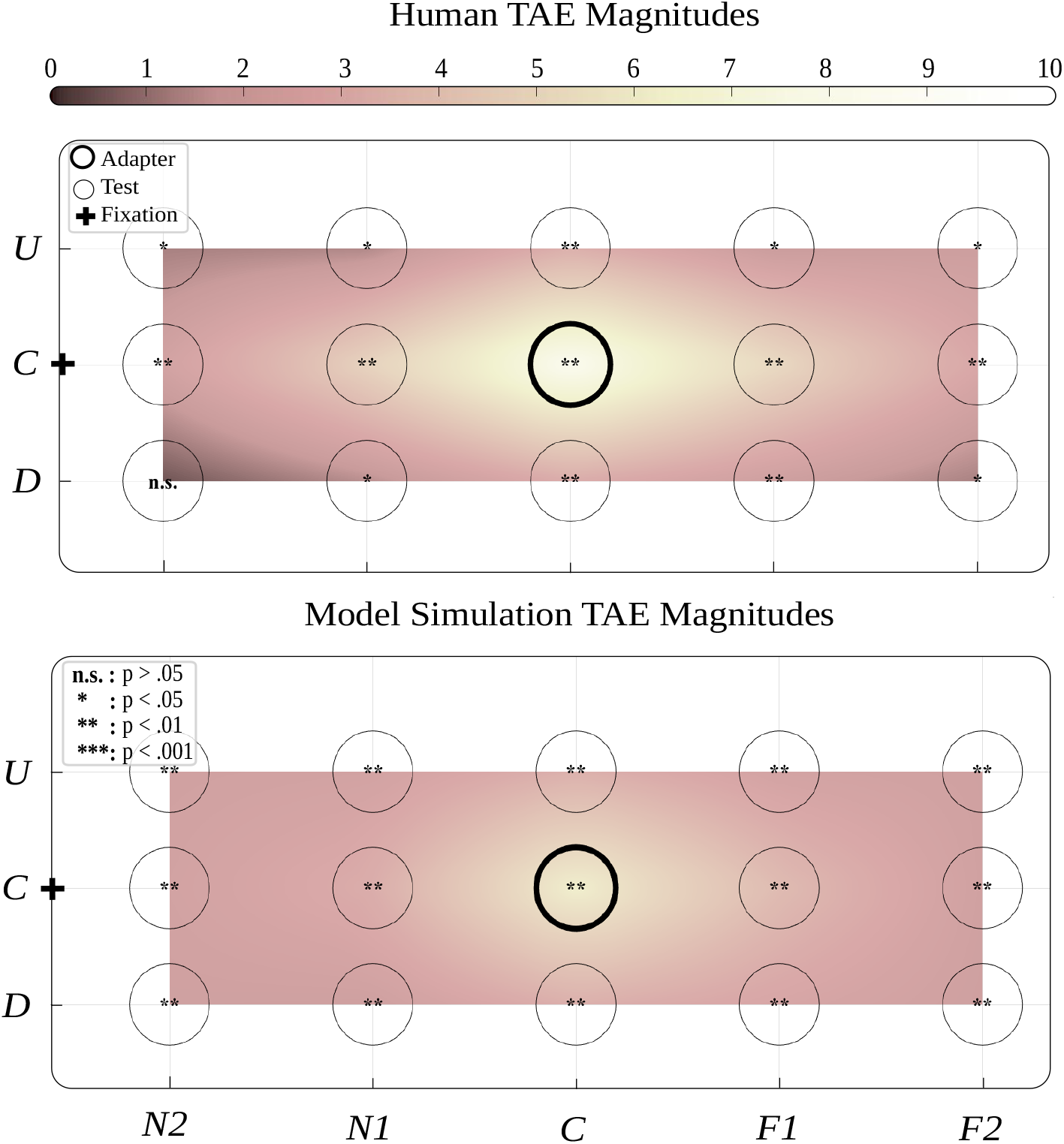
The TAE magnitudes calculated for human behavioral experiment (top panel) and model simulation (bottom panel) shown as a heatmap. The human map was generated by averaging across the two visual hemifields because there was no significant difference between the hemifields. The TAE magnitudes for the positions between the test positions were calculated by natural neighbor interpolation. As shown in the top color bar, brighter colors indicate higher TAE magnitudes. The cross sign shows the fixation point (FP) location, and circles represent the test positions. Naming convention for the test positions is the same as in Figure 2. The bold circle located at the center row and center column (*C*_*c*_) indicates the location where adapter patch was presented. Both maps demonstrate that TAE is a non-local aftereffect and spreads across the visual hemifield. Star signs: FDR correct significance levels.

Furthermore, the spread of TAE depended on the location of the test patch. Specifically, within-subject effects of the mixed ANOVA revealed a significant interaction of test rows and columns (Sphericity was corrected by Greenhouse-Geisser correction *F* (4, 61) = 41.33, *p* = 0.01), as shown in Figure 5. As can be observed with a visual inspection of the figure, the TAE magnitude was significantly higher at the center row where the adapter and FP were located compared to other rows (FDR corrected *ps <* 0.005) for all test columns except *N2* and *F2*. Pairwise comparisons of the test columns showed significantly higher TAE magnitude at the Center column where the adapter was located compared to all other columns (FDR corrected *ps* ≤ 0.004). The TAE magnitude decreased with the distance between the test and the adapter. This decrease was proportional to the absolute value of the distance and did not depend on the direction (no significant difference between *N2* and *F2* columns (FDR corrected *p* = 0.1), and between *N1* and *F1* columns (FDR corrected *p* = 0.1), and between Up and Down rows (FDR corrected *p* = 0.33)).

**Figure 5:**
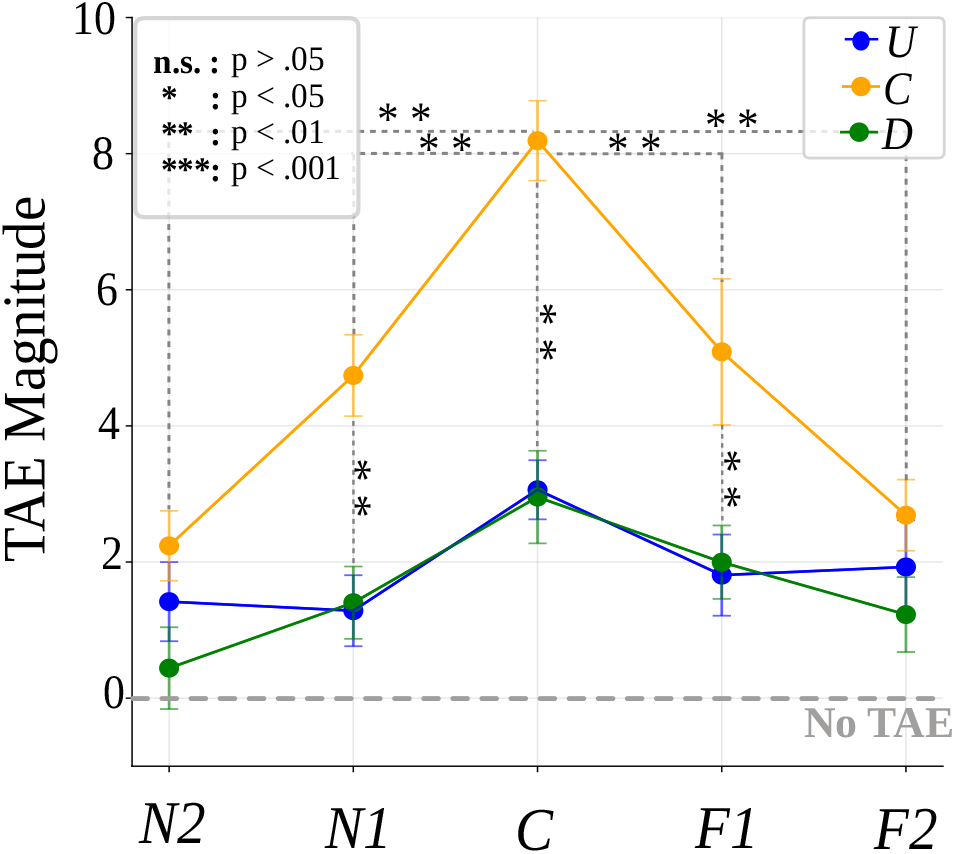
Human TAE magnitudes are re-plotted. The x-axis represents the test columns, and color-coded lines represent the test rows (the naming convention is the same as in Figure 2). The within-subject effects of the mixed ANOVA revealed a significant interaction of test rows and columns, showing that the spread of the TAE depends on the location of the test grating. Star signs: FDR corrected significance levels. Error bars: standard error of the mean.

## Computational Model

Next, in order to establish a link between the behavioral results and possible underlying neuronal mechanisms, we have developed and tested a biologically plausible toy model (publicly available, see Data and Code Availability Statement section). The model had an encoding and a decoding component (layer). The encoding component was composed of populations of orientation-tuned units on a grid covering the visual hemifield. The decoder component estimated the orientation and tilt based on the input it received from the encoding component. While calculating the responses of the orientation-tuned units in the encoding component, we used both shift and suppression mechanisms of adaptation (Jin et al., 2005). Accordingly, our model assumed that, after adaptation, unit responses could decrease (suppression) and their preference could shift away from the adapter orientation as shown in Figure 1. For the model fits to observer data, we optimized the shift and suppression parameters, as well as the RF sizes using a machine learning procedure. We were particularly interested in the optimized RF sizes, because they could potentially inform us about possible cortical origins of the spread of TAE.

### Model Architecture & Optimization

We simulated the encoder unit responses on a 20 × 20 visual grid across a hemifield, where lattice points were separated by 1° of visual angle. At each lattice point we positioned a population of orientation-tuned units (*N* = 180) modeled using a Gaussian-shaped function where the maximum response was equal to 1, and half-width at halfheight was equal to 30° (Westrick, Heeger, & Landy, 2016). *The preferred orientations of these units were spaced by 1*° polar angle. Inspired by the term ‘hyper-column’ in primary visual cortex (V1), we will refer to these multi-unit structures at each lattice point as *hyper-units*.

The response of an encoder unit with receptive field center *x, y*, and preferred orientation *ψ* is computed as a weighted product of its tuning curve and RF profile

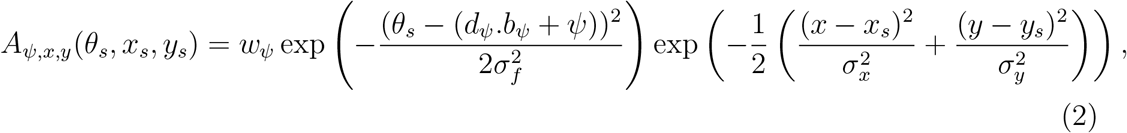

where *x*_*s*_, *y*_*s*_ and *θ*_*s*_ define the stimulus position and orientation, *σ*_*f*_ defines the width of the tuning curve, *σ*_*x*_ and *σ*_*y*_ are the size parameters for the two dimensional RF profile, and *w*_*ψ*_ and *b*_*ψ*_ are the parameters defining the amount of suppression and shift of the tuning curve of that unit after adaptation (we have dropped the indices *x, y* for clarity). The direction of the tuning curve shift was determined by *d*_*ψ*_ = sgn(*ψ* − *θ*_*a*_) to ensure that the shift was always away from the adapted orientation *θ*_*a*_ (sgn(), sign function).

To model the suppression and shift, we first computed the unit responses during the adapter period using Eq.2 with *w*_*ψ*_ = 1, *b*_*ψ*_ = 0, and stimulus variables *θ*_*a*_, *x*_*a*_, *y*_*a*_. We assumed that the adaptation of a unit is related to its activation during the adapter period, and modeled this relation using power functions

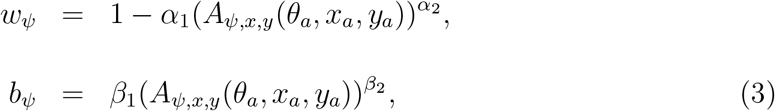

with the constrains *w*_*ψ*_ ∈ [0, 1] and *b*_*ψ*_ ≥0. We also assumed *w*_*ψ*_ = 1 and *b*_*ψ*_ = 0 during the no-adaptation trials.

Based on the responses from the encoder layer, the decoder estimated the orientation of the input stimulus 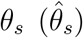 for stimulus location (*x*_*s*_, *y*_*s*_) using the winner-take-all method with a suitable differentiable approximation to the argmax function

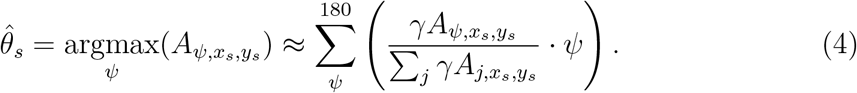

Next, the decoder returned a probability that the input stimulus is tilted clockwise as

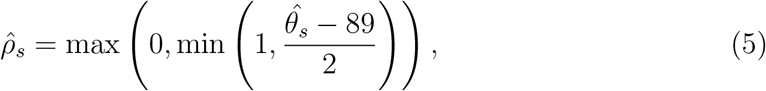

where 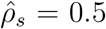 when the predicted stimulus orientation is vertical 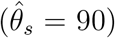 Table 1 summarizes all the variables and optimized parameters in our model.

**Table 1:** Variables and optimized parameters used in the computational model.

| Symbol | Description |
| --- | --- |
| $\theta_a, x_a, y_a$ | orientation & position of the adapter grating |
| $\theta_s, x_s, y_s$ | orientation & position of the test grating |
| $\psi, x, y$ | preferred orientation & RF center of a simulated encoder unit |
| $\sigma_f$ | width of the orientation tuning curve of an encoder unit |
| $\sigma_x, \sigma_y$ | two-dimensional RF size parameters of an encoder unit, optimized |
| $A_{\psi, x, y}(\theta_s, x_s, y_s)$ | activation of an encoder unit |
| $d_\psi, b_\psi$ | direction and magnitude of tuning curve shift of the encoder unit<br>with preferred orientation $\psi$ |
| $w_\psi$ | suppression parameter of an encoder unit with preferred orientation $\psi$ |
| $\beta_1, \beta_2$ | parameters of $b_\psi$ power function, optimized |
| $\alpha_1, \alpha_2$ | parameters of $w_\psi$ power function, optimized |
| $\hat{\theta}_s$ | predicted orientation of the test grating by the decoder |
| $\hat{\rho}_s$ | probability that test grating is tilted clockwise as reported by the decoder |

We implemented and optimized the model using the PyTorch library in Python (Paszke et al., 2019). Training was done on randomly selected 80% of the adaptation condition data. Testing the model performance was done on the remaining 20% of the adaptation condition data, as well as on randomly selected 20% of the no-adaptation condition data. We trained the model separately for each participant and optimized *b*_*ψ*_, *w*_*ψ*_, *σ*_*x*_, and *σ*_*y*_ parameters. Optimization was performed using the adaptive moment estimation (Adam) optimization (Kingma & Ba, 2014). The model architecture was the same for adaptation and no-adaptation conditions, but because no suppression and shift are expected on the no-adaptation condition, we set *b*_*ψ*_ = 0 and *w*_*ψ*_ = 1, and used the *σ*_*x*_ and *σ*_*y*_ parameters optimized for the adaptation condition. Therefore, we did not perform training for no-adaptation condition and this simulation served as a control for our model’s performance.

### Model Performance & Simulations

To measure the model’s performance, we tested the model on the test sets for both adaptation and no-adaptation conditions. We calculated the accuracy as the percentage of correctly predicted tilt directions and used this as a metric of model performance evaluation.

Next, using the optimized parameters per participant, we made our model run the same behavioral experiment for adaptation and no-adaptation conditions. Based on the simulated responses, we first obtained the TAE magnitudes via the same method used in the behavioral experiment. Then, we performed a two-tailed one sample t-test to test whether TAE magnitudes are significantly different from zero at each location (multiple comparisons were corrected by FDR correction), and formed the same TAE-magnitude heat-map as in the behavioral experiment.

### Model Results

The model could predict the human behavioral data substantially above chance level (50%) for adaptation and no-adaptation conditions at all test locations (Figure 6). The average accuracy for the adaptation condition was 76.9% (*SD* = 0.13), and the average accuracy for the no-adaptation condition was 66.7% (*SD* = 0.19). Furthermore, a visual inspection of model simulation and human TAE magnitude maps in Figure 4 reveals a closely matching pattern. These results show that the model can successfully predict the human data.

**Figure 6:**
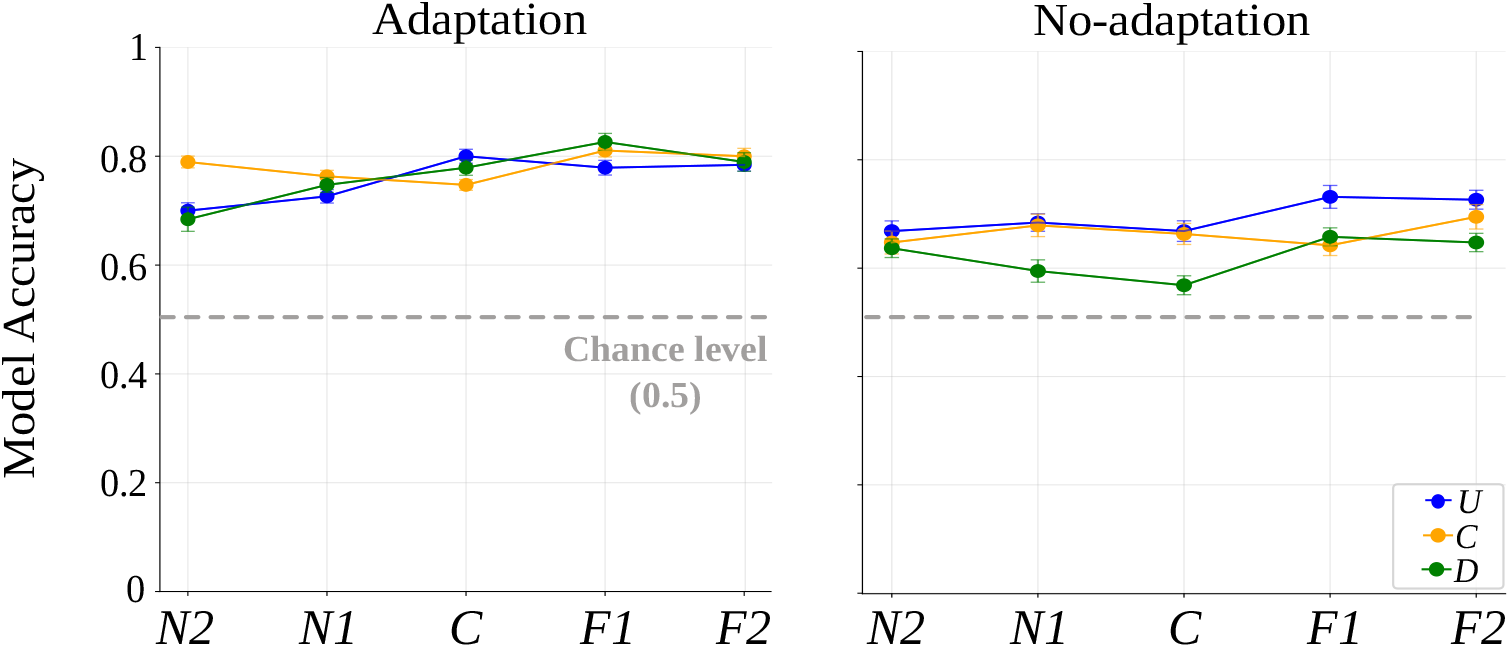
Model accuracy for adaptation (left panel) and no-adaptation (right panel) conditions. The x-axis and color-coded lines represent test columns and rows, respectively (the naming convention is the same as in Figure 2). First, we fit the model to each participant’s data. Then, we used the trained model to predict unseen data of the participant and calculated the accuracy. Finally, we averaged the obtained accuracy values across participants. Error bars: standard error of the mean.

Next, to validate that the model behaves reasonably, we evaluated the optimized suppression and shift values. Table 2 reports the optimized parameters of the power functions (Eq. 3), and Figure 7 shows the suppression and shift values derived from those parameters. Consistent with the behavioral results, and as expected based on the findings in literature, the model predicted stronger suppression and larger shift for units whose RF centers are closer to the adapter position, and whose preferred orientations are closer to the adapter orientation. Suppression and shift decreased as the distance between the unit RF center and the adapter position increased, and as the difference between the preferred orientation of the unit and the adapter orientation increased. These results show that the internal workings of the model is consistent with the behavioral results and findings in literature.

**Table 2:** Optimized parameters of the power functions determining suppression and shift (*w*_*ψ*_ and *b*_*ψ*_, see Eq. 3). The model was first fit to each participants’ data and then the results were averaged across participants.

| $\alpha_1$ | $\alpha_2$ | $\beta_1$ | $\beta_2$ |
| --- | --- | --- | --- |
| 0.67 | 0.29 | 0.74 | 0.17 |

**Figure 7:**
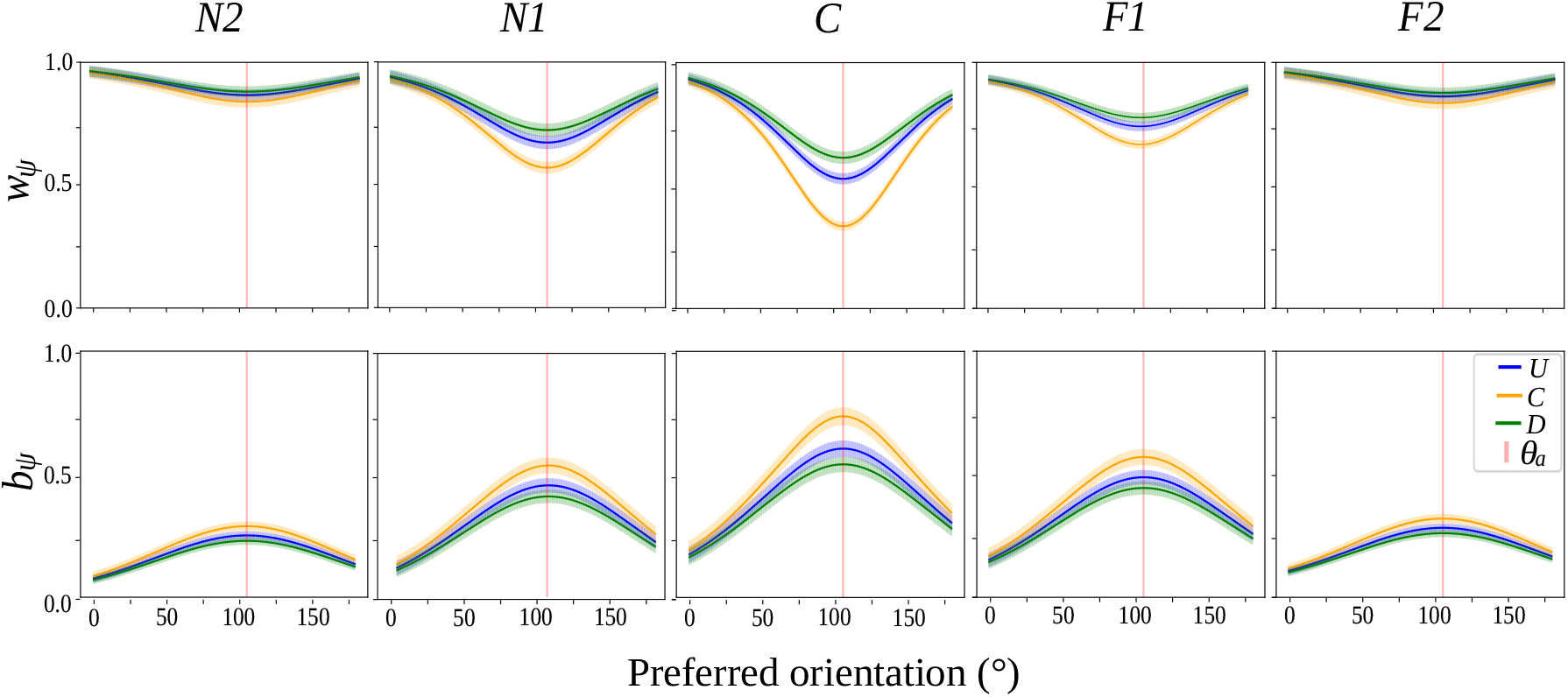
Suppression, *w*_*ψ*_, and shift, *b*_*ψ*_, values predicted by the model. Smaller *w*_*ψ*_ values mean stronger suppression, larger *b*_*ψ*_ mean larger shift. Panels and color-coded solid lines indicate the RF center positions and the x-axes indicate the preferred orientations of the encoder units (panels: horizontal positions; color-coded lines: vertical positions). The thin vertical red lines mark the adapter orientation (*θ*_*a*_). Transparent thickness of the solid lines represents the standard error of the mean. Stronger suppression and shift are observed at units with RF centers and preferred orientations closer to the adapter position and orientation.

Finally, we evaluated the RF size parameters *σ*_*x*_ and *σ*_*y*_. Table 3 reports their optimized values. This analysis was particularly important because optimized RF size parameters can potentially provide clues about the cortical origins of the behavioral effect. Overall *σ*_*x*_ and *σ*_*y*_ values were around 3 degrees (mean *σ*_*x*_: 3.09, *SD* = 0.12; mean *σ*_*y*_: 3.06, *SD* = 0.11). Within each hyper-unit *σ*_*x*_ and *σ*_*y*_ values did not differ much from each other, which means that the RFs were approximately isotropic. Further, there was no substantial difference between the optimized values for different test locations, which suggests that the modeled RF sizes did not vary with eccentricity (spatially invariant). We discuss possible implications of these findings in the next section.

**Table 3:** Optimized values of two-dimensional RF size parameters for all test locations. Same as in Table 2, the values were obtained by averaging the optimized RF size parameters across participants.

| $(\sigma_x, \sigma_y)$ | $N2$ | $N1$ | $C$ | $F1$ | $F2$ |
| --- | --- | --- | --- | --- | --- |
| $U$ | (3, 3) | (3.3, 3.3) | (3.15, 3) | (3.12, 3.12) | (3, 3) |
| $C$ | (3, 3) | (2.95, 2.95) | (3.01, 3) | (3.05, 3.05) | (3, 3) |
| $D$ | (3, 3) | (3.18, 3.18) | (3.25, 3) | (3.27, 3.27) | (3, 3) |

## Discussion

Here, we studied whether the TAE is localized at the position of the adapter stimulus or it spreads across the visual field. Consistent with the previous literature, our behavioral results showed that prolonged view of an oriented Gabor patch (adapter) led to changes in the perceived orientation of the subsequently presented test patch at the location of the adapter. Specifically, the perceived orientation of the test patch was repelled away from the orientation of the adapter. But most importantly, our results showed that TAE is not limited to the adapter location; instead it spreads across the visual hemifield. The spread was systematic: TAE was highest at the adapter location, and decreased with the distance between the adapter and the test in a nearly isotropic manner.

Previous models based on the local TAE results supported the idea that the effect originates in V1 through feedforward mechanisms (e.g. Jin et al., 2005). Could a similar V1 model explain our results here? Or does the spread of the effect require the involvement of other mechanisms? Conceptually, in a feedforward mechanism, the spread of the effect would require involvement of neurons with relatively large receptive fields (RFs), which can integrate information across the visual hemifield. To test this, we next developed a computational model and estimated the RF sizes of hypothetical neurons whose activity could predict the specifics of the spread we observed in our behavioral data.

The computational model could indeed successfully predict the empirical TAE magnitudes at all tested locations in the visual hemifield with high accuracy. Further, the model captured the pattern of the empirical TAE magnitudes across the hemifield. Specifically, it predicted the highest TAE at the location of the adapter, and the simulated TAE magnitudes decreased nearly isotropically as the distance between the adapter and test increased. As we have expected, the optimized RF sizes were relatively large (approximately 3 degrees, on average), and they did not change substantially with eccentricity. Consequently, an ideal candidate visual area whose activity gives rise to the spread of TAE across the hemifield would be the one with relatively large RFs while having selectivity to orientation.

Aforementioned qualities are commonly associated with mid-level visual areas; for example the visual area V4. V4 is situated along the ventral pathway, and its characteristics include selectivity to orientation and color (Desimone & Schein, 1987; Roe et al., 2012). Its activity is linked to surface perception and completion (Bouvier, Cardinal, & Engel, 2008; Pan et al., 2012), integration of spatial information across the visual space, and perception of illusory contours (De Weerd, Desimone, & Ungerleider, 1996). RF sizes of V4 neurons were found to be 4 to 7 times larger than those of V1 neurons in macaque (Desimone & Schein, 1987). Similarly, a human functional magnetic resonance imaging (fMRI) study found that V4 population receptive fields are larger than those in V1, and importantly, their sizes do not change substantially with eccentricity (van Dijk, de Haas, Moutsiana, & Schwarzkopf, 2016). These RF characteristics are in line with our model predictions. Furthermore, V4 is extensively interconnected with other visual areas. It receives direct inputs from V1 (Nakamura, Gattass, Desimone, & Ungerleider, 1993), and it is connected to higher-level temporal areas such as TE and TEO, which suggests that it plays a role in object recognition. It is also connected to higher-level dorsal areas such as DP, VIP LIP, PIP, and MST, which suggests that it plays a role in spatial vision and attention, as well (Baizer, Ungerleider, & Desimone, 1991; Ungerleider, Galkin, Desimone, & Gattass, 2008). Thus, we contend that a midlevel visual area, such as V4, appears to be a plausible candidate to be the origin of the spread of TAE across the visual field. However, the exact origin of the effect would require further neuronal investigation.

It should be noted that here we proposed a simple feedforward model. Alternative models incorporating feedback can also be tested. For example V1 activity modulated by feedback from higher-level visual areas may have a causal relation with the spread of the effect. Therefore, future studies testing such models in conjunction with neuronal activity in V4 and other visual areas would be required for a better understanding of the origin of the spread of figural aftereffects including TAE and the size aftereffect (Altan & Boyaci, 2020).

### Possible role of spatial attention

Encoding the spatial location of a stimulus is argued to be a primary or mandatory process. For example, it was shown that attending to a stimulus feature, with the location being irrelevant, was automatically accompanied by attention to the spatial location that the stimulus occupies (Tsal & Lavie, 1993). Therefore, in our experimental paradigm, adapting to a tilted contour that always appears at the same location within the visual hemifield could automatically trigger spatial attention to that location, and enhance the neuronal responses (Gandhi, Heeger, & Boynton, 1999; Treue & Trujillo, 1999). This could play a role for the highest TAE observed at the adapted location. This prediction is consistent with previous studies that demonstrated higher magnitudes of perceptual aftereffects at the attended spatial locations (e.g., Yeh, Chen, De Valois, De Valois, et al., 1996). Even though our model integrates some attentional mechanism by the winner-take-all strategy (Lee, Itti, Koch, & Braun, 1999) to predict the perceived tilt, attention could be explicitly manipulated in future behavioral experiments and its effects could be more explicitly incorporated into the current model architecture to test these hypotheses further (e.g., the normalization model of attention Reynolds & Heeger, 2009).

### Potential applications for visual rehabilitation, and implications for perceptual learning

Developing rehabilitation techniques to restore, recover or improve the vision of patients with neural damage is an integral part of vision research. Findings from the literature show that vision could be restored with appropriate perceptual training methods. However, this recovery is often believed to be limited to the trained location and not transfer to other locations in the visual field (for example in blindsight, Cowey & Stoerig, 1991). Our findings, on the other hand, could encourage work on development of more efficient training techniques. Specifically, neurons at higher-level visual areas with large RFs could respond to the training stimulus that is presented at different parts of the visual field. Accordingly, training with a stimulus that is preferred by neurons in those visual areas could lead to visual improvements that are not specific to the exact location of that stimulus. Indeed, training blindsight patients with a stimulus targeting higher-level visual areas (e.g., a complex optic flow motion stimulus) lead to improvements in untrained locations (Awada, Bakhtiari, Legault, Odier, & Pack, 2022). Further in line with this idea, recent studies in perceptual learning literature provide evidence that learning transfers to other locations with training paradigms that leverage the involvement of higher-level visual areas (Bakhtiari, Awada, & Pack, 2020). These training effects could be explained in part with our findings here.

### Implications for computer vision

Recently artificial neural networks (ANNs) have been remarkably successful in computer vision tasks such as object recognition and image segmentation. Yet, these networks do not usually employ temporal and spatial integration together. In particular, sequential training of ANNs through time with a visual feature, does not lead to altered perceptions at other locations. Our results here, however, show that such integration processes happen in human visual system, which could be important for more efficient computational processing of the visual information. Therefore, our findings here could have implications for ANNs to achieve more efficient and human-like computer vision systems.

## Conclusion

To conclude, we have found that prolonged exposure to a tilted Gabor patch affects the perceived orientation of subsequently presented stimuli not only at the location of the adapter, but also in other parts of the visual field. Moreover, a simple feedforward computational model was able to predict the systematic pattern in the human data. Importantly the optimized RF sizes in the model suggest that the spread of the effect could be originating in a mid-level visual area, such as V4. These results may also have implications for developing more efficient visual rehabilitation methods and computer vision algorithms.

## Data and Code Availability Statement

The code for the experimental paradigm and computational model together with the anonymized data of this study is publicly available under the GitHub page https://github.com/tugcegurbuz/TAE-spreads-across-the-visual-field.

## CRediT authorship contribution statement

**Busra Tugce Gurbuz:** Conceptualization, Methodology, Software, Data Collection, Analysis, Visualization, Writing - original draft. **Huseyin Boyaci:** Conceptualization, Supervision, Writing - review & editing.

## Declaration of Competing Interest

The authors declare that they have no known competing financial interests or personal relationships that could have appeared to influence the work reported in this paper.

## Acknowledgments

This research did not receive any specific grant from funding agencies in the public, commercial, or not-for-profit sectors.

## Notes

https://github.com/tugcegurbuz/TAE-spreads-across-the-visual-field

### Competing Interest Statement

The authors have declared no competing interest.

